# Natural influent bioaugmentation of activated sludge water resource recovery systems: implications for low temperature nitrification and heterotrophic population structures

**DOI:** 10.1101/2021.10.03.462950

**Authors:** Shameem Jauffur, Zeinab Bakhshi, Dominic Frigon

## Abstract

This work aimed at demonstrating the natural bioaugmentation of biological activated sludge systems with nitrifying biomass from influent wastewater in lab-scale reactors. Three sequencing batch reactors (SBR) were fed with sterile synthetic wastewater. While nitrification was complete at a temperature of 8 °C and a SRT of 20 days, it failed when the temperature was lowered to 5 °C, and the SRT decreased to 7 days. In the test period, the sterile synthetic wastewater fed to the Test Reactor was supplemented by influent solids harvested at a full-scale treatment facility at a total suspended solids concentration of 100 mg/L, which corresponded to approximately 5 mg-COD/L of nitrifying biomass. Upon this addition, nitrification was restores. Subsequent halting the supply of influent solids to the Test Reactor led a rapid failure of nitrification and washout of nitrifiers from the SBR. Reproducibility was demonstrated by switching the feed composition between the Test and Negative control reactors. PCR-based amplicon sequencing analyses targeting the *amoA*, and *nxrB* genes of the *Nitrospira* genus have shown that the influent wastewater governed the structure and composition of the activated sludge nitrifying populations. The most abundant ammonia-oxidizing bacteria (AOB) and *Nitrospira*-related nitrite-oxidizing bacteria (NOB) in the influent seeds occurred as the most dominant ones in the activated sludge. This pattern was observed even when the influent seeds varied over time. The heterotrophic populations were less affected by the influent seeds with the activated sludge and raw sewage showing distinct microbial populations based on principal coordinate analysis (PCoA). However, the immigrant populations appeared to modulate the structure of the activated sludge heterotrophic communities to some extent. These findings provide concrete evidence of the presence of active nitrifiers in raw wastewater capable of supporting nitrification in an otherwise non-conducive environment. This may have important implications on process design, operation and optimization of wastewater treatment systems.

**Highlights:**

- Lab-scale reactors fed sterile synthetic wastewater at low temperatures and SRTs.
- Nitrification failed when conditions were adjusted to 5 °C and a SRT of 7 days.
- Nitrification restored by addition of real wastewater influent solids.
- Nitrifiers in solids from sewers naturally bioaugment activated sludge systems.
- Activated sludge models should consider the immigration of nitrifiers with influent.

**Graphical abstract:** 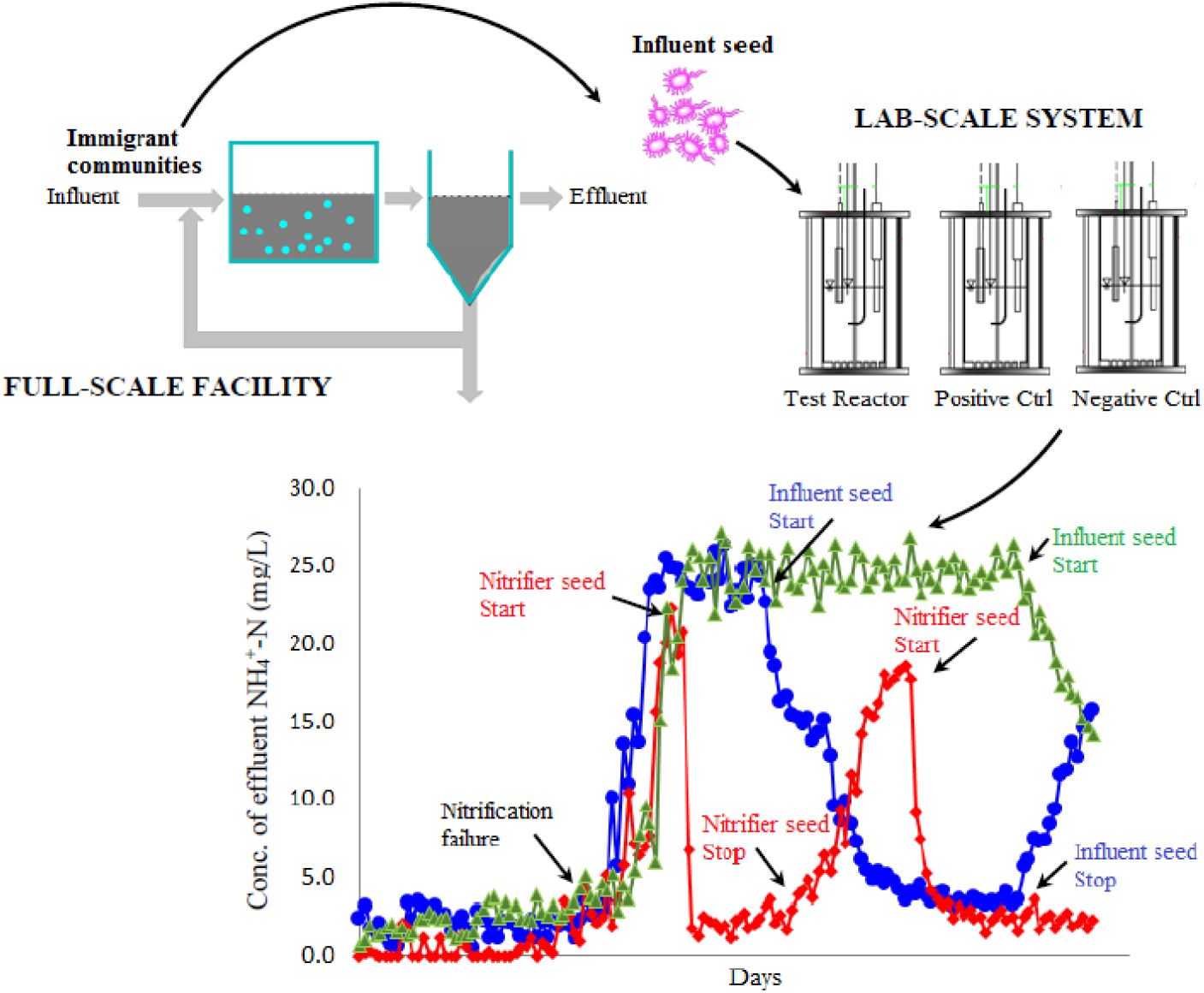

## 1. Introduction

Although bacterial biomass is commonly quantified in activated sludge for wastewater treatment process design and operation (Ekama 1984, Henze 2000), it is rarely quantified in raw wastewater, and is generally neglected in mass balances due to low concentrations (Foladori et al. 2010). Even best modeling practices adopted by the International Water Association (IWA) consensus Activated Sludge Models (ASMs) assume no active biomass in influent wastewaters. According to Grady Jr et al. (2011), two significant effects may result if active biomass is present in influent wastewaters. It will reduce the minimum substrate concentration attainable (*S*_s,min_), and prevent washout even at low solids retention time (SRT). This underlies the concept of bioaugmentation of biomass in activated sludge systems that enhances process performance.

Recently, the presence of nitrifiers has been reported in raw wastewaters reaching full-scale wastewater treatment facilities (Jauffur et al. 2014, Saunders et al. 2015). The nitrifiers detected in the wastewater were shown to be alive with the potential of being fully induced metabolically within a few hours (Jauffur et al. 2014). Nitrifiers from influent wastewaters may be adsorbed to activated flocs and be retained in the bioreactor to provide additional nitrification activity. Moreover, it has been observed that dominant nitrifying operational taxonomic units (OTUs) identified in municipal influent wastewaters and corresponding activated sludge mixed liquors of full-scale Water Resource Recovery Facilities (WRRFs) were the same, suggesting that natural seeding of nitrifiers occur at full-scale wastewater treatment facilities (Jauffur et al. 2018, Unpublished data). This may be of significant importance especially in cold conditions where nitrification may be compromised (Van Dyke et al. 2003). Further research is necessary to understand the importance of influent wastewater in supplying indigenous nitrifying bacteria to activated sludge systems. It is believed that these nitrifiers are conveyed to combined sewer systems by wastewaters and rainwater inputs, as well as infiltration from soil water into the sewer pipes. Sewer infiltration has been shown to be particularly prevalent in ageing and leaky infrastructures (Kuroda et al. 2012). Such infiltration fluxes can increase during precipitation events thereby increasing the water levels and hydraulic potential of sewer systems (Musolff et al. 2010). Nitrifiers, which are abundant in soils where they are responsible for nitrogen cycling and regulating N-availability to vegetation (Tsiknia et al. 2014), may thus infiltrate and colonize sewer systems.

Influent wastewaters also contain a wide diversity of heterotrophic bacterial communities resulting from inputs of human fecal matter and water infiltration, or from growth of specific microbes in the sewer system (McLellan et al. 2010). A fresh sewage usually contains a limited supply of oxygen and most of the internal sewer system can be considered to be under sulphate-reducing anaerobic conditions (Guisasola et al. 2008), thereby favoring the growth of anaerobic fermenters (Nielsen 1991). However, it is possible to achieve partial oxidation of organic substrates by resident microorganisms during sewage conveyance. Gravity sewers have been shown to abate the concentration of organic matter when the dissolved oxygen (DO) level in the water phase is greater than 1 mg/L with active bacteria growing in the sediments and as biofilms on the side walls of sewers (Chen et al. 2003). According to Lemmer et al. (1994), biofilms developed on the inner wall of sewer pipes, can have activities one or two magnitudes higher than that of activated sludge microorganisms. Erosion, sloughing and sewer conveyance of biomass by wastewaters lead to immigration of microbial communities, which contribute to observed bacterial diversities in activated sludge. Thirty-five percent of heterotrophic bacterial OTUs identified in activated sludge was found to come from influent wastewaters in Danish WRRFs (Saunders et al. 2015).

The aim of this study was to reproduce in the laboratory, the natural bioaugmentation of nitrifiers by influent wastewaters observed at full-scale WRRFs. The laboratory reactors fed with sterile culture medium did not support nitrification under conditions of low operational temperature and SRT. However, ASM model simulations with typical default parameters suggest that nitrification should have occurred under these conditions. Thus, this study evaluated the importance of influent nitrifiers to support nitrification in activated sludge bioreactors. In addition, the impact of influent wastewater seeding on heterotrophic microbial community composition in activated sludge was also assessed.

## 2. Material and methods

### 2.1. Sequencing batch reactor (SBR) set-up and operation

Three 4-L cylindrical double-wall jacketed laboratory-scale sequencing batch reactors (SBRs) made up of non-reactive Plexiglas were set-up with an actual working volume of 2 L. A schematic layout of the SBRs is shown in Fig. 1. The temperature of the reactors was controlled using thermostatic water circulators (IsoTherm model 250LC, Fisher Scientific, MA). Influent, effluent and waste activated sludge (WAS) were pumped by means of Masterflex peristaltic pumps (Model SI-77911-20, Cole Parmer, USA). The operation of the SBR systems was automated using the Apex AquaController (model APEXLSYS, Neptune Systems, San Jose, CA), which also controlled the pH to 7.45 by titrating 0.25 M NaOH. The SBRs were aerated at an airflow rate of 2 L/min using compressed air, which was filtered and introduced through diffusion plates at the bottom of the reactors. During start-up, the reactors were inoculated with activated sludge mixed liquor suspended solids (MLSS) collected from the Régie d’Assainissement des Eaux du Bassin La Prairie (RAEBL), a full-scale biological WRRF located near Montreal (Canada). The reactors were operated with a 6-h cycle. At the beginning of the cycle, aeration was started for 10 min prior to pumping 1 L of feed into each reactor for 5 min, followed by an aeration phase of 5 h. Prior to the end of the aeration phase, a specific volume of activated sludge was wasted to keep the SRT at a constant level. This was followed by a settling phase of 0.75 h after which 1 L of supernatant was removed resulting in a hydraulic retention time (HRT) of 0.5 day.

**Fig. 1.**
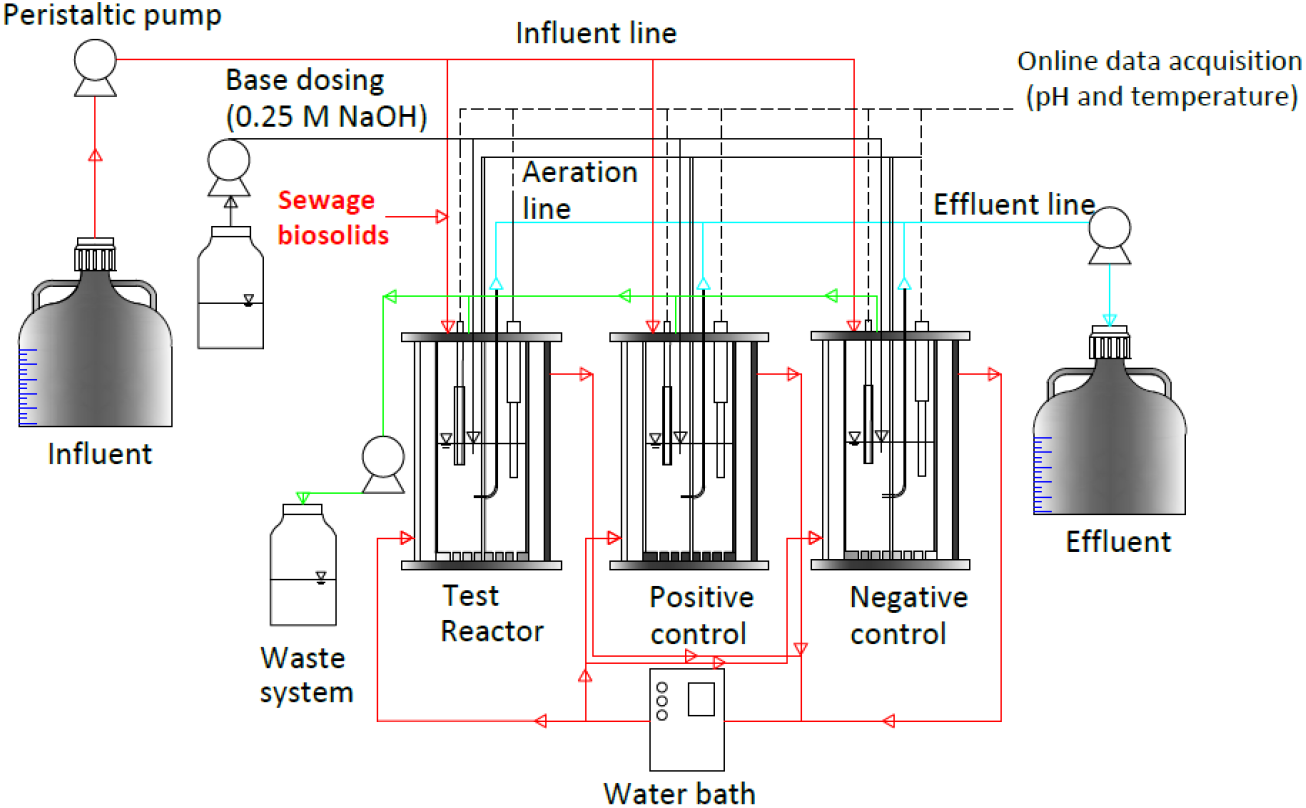
Schematic overview of sequencing batch reactors (SBRs). Set-up comprised of an influent line feeding synthetic wastewater (*Syntho*) to three SBRs (Test Reactor, Positive Control and Negative Control reactors) and an effluent line drawing treated effluent out of the system. Each treatment cycle (fill, react, settle, decant and idle) lasted for 6 h.

The three SBRs were fed with “*Syntho*”, a synthetic feed developed by Boeije et al. (1999), which mimics the average composition and quality of domestic wastewater. The composition of 1 L of synthetic feed was as follows: 15 mg peptone, 120 mg CH_3_COONa, 15 mg dry meat extract, 40 mL glycerol, 50 mg starch, 120 mg low fat milk powder, 75 mg urea, 9 mg uric acid, 11 mg NH_4_Cl, 25 mg MgHPO_4_.3H_2_O, 20 mg K_3_PO_4_.H_2_O, 10 mg sodium dodecyl sulfate (SDS), 10 mL Synapol and 10 mg diatomaceous earth. A mixture of trace elements based on typical concentrations found in sewage was added to the “*Syntho*” (Bollmann et al. 2011) and comprised of the following in 1 L: 4,292 mg NaEDTA, 2,780 mg FeSO_4_.7H_2_O, 99 mg MnCl_2_.4H_2_O, 24 mg NiCl_2_.6H_2_O, 24 mg CoCl_2_.6H_2_O, 13.4 mg CuCl_2_, 143 mg ZnSO_4_.7H_2_O, 24 mg Na_2_MoO_4_.2H_2_O, 23.2 mg WO_3_ and 62 mg H_3_BO_3_. The feed (without the trace element solution) was autoclaved at 121 °C for 1 h and allowed to cool down prior to feeding the reactors. The COD and TKN of the formulated influent recipe was approximately 550 mg/L and 35 mg-N/L, respectively.

An additional SBR operated at a SRT of 15 days and HRT of 0.5 day, was used to enrich nitrifying biomass at 5 °C by feeding it with a recipe based on Bollmann et al. (2011). The mineral medium consisted of the following in 1 L: 660.7 mg (NH_4_)_2_SO_4_, 840 mg NaHCO_3_, 585 mg NaCl, 75 mg KCl, 147 mg CaCl_2_.2H_2_O, 49 mg MgSO_4_.7H_2_O, 1360.9 mg KH_2_PO_4_, 57.2 g HEPES buffer, 240 mg diatomaceous earth and trace element with the same chemical composition used to prepare the “*Syntho*” organic feed. Stock solutions were autoclaved at 121 °C for 1 h before use to prevent bacterial or fungal growth during storage. The pH of the mineral medium was adjusted to 7.8 using 1 M NaOH. The reactor was aerated with filtered compressed air enriched with 0.4% (v/v) CO_2_ gas to support the autotrophic growth of nitrifiers (Denecke and Liebig 2003). The pH of the reactor was maintained at 7.45 by adding 0.25 M NaOH using an automatic base dosing system.

### 2.2. Natural influent seeding experiments

The SBRs were operated at a temperature of 8 °C and a SRT of 20 days for the initial 130 days (Phase I in Fig. 2). Prior to the start of the seeding experiment, the temperature of the three SBRs was lowered to 5 °C, and the SRT was decreased to 7 days to cause nitrification failure in all the three reactors (Phase II in Fig. 2). Following nitrifier washout and failure of the reactors to nitrify ammonium, the seeding experiments were started to find out whether seeding with municipal influent solids could re-establish nitrification and stabilize the bioreactors. One of the SBRs (Test Reactor) was seeded with influent solids harvested from the influent of the RAEBL full-scale WRRF by weekly centrifuging 10 L of a 24-h composite influent sample at 4,000×g using the IEC-Centra-8 centrifuge (Thermo Scientific, USA). The total suspended solids (TSS) concentration was adjusted to 100 mg/L, which corresponded to a nitrifier population concentration of approximately 5 mg-COD_biomass_/L (representing ~1.5% of the total COD), a level similar to what was observed at the full-scale RAEBL WRRF based on respirometric assays (Jauffur et al. 2014). Another SBR, referred to as the “Positive Control” was seeded with cold-adapted enriched nitrifying culture at a similar rate. The third SBR was used as a “Negative Control” reactor and was not seeded.

**Fig. 2.**
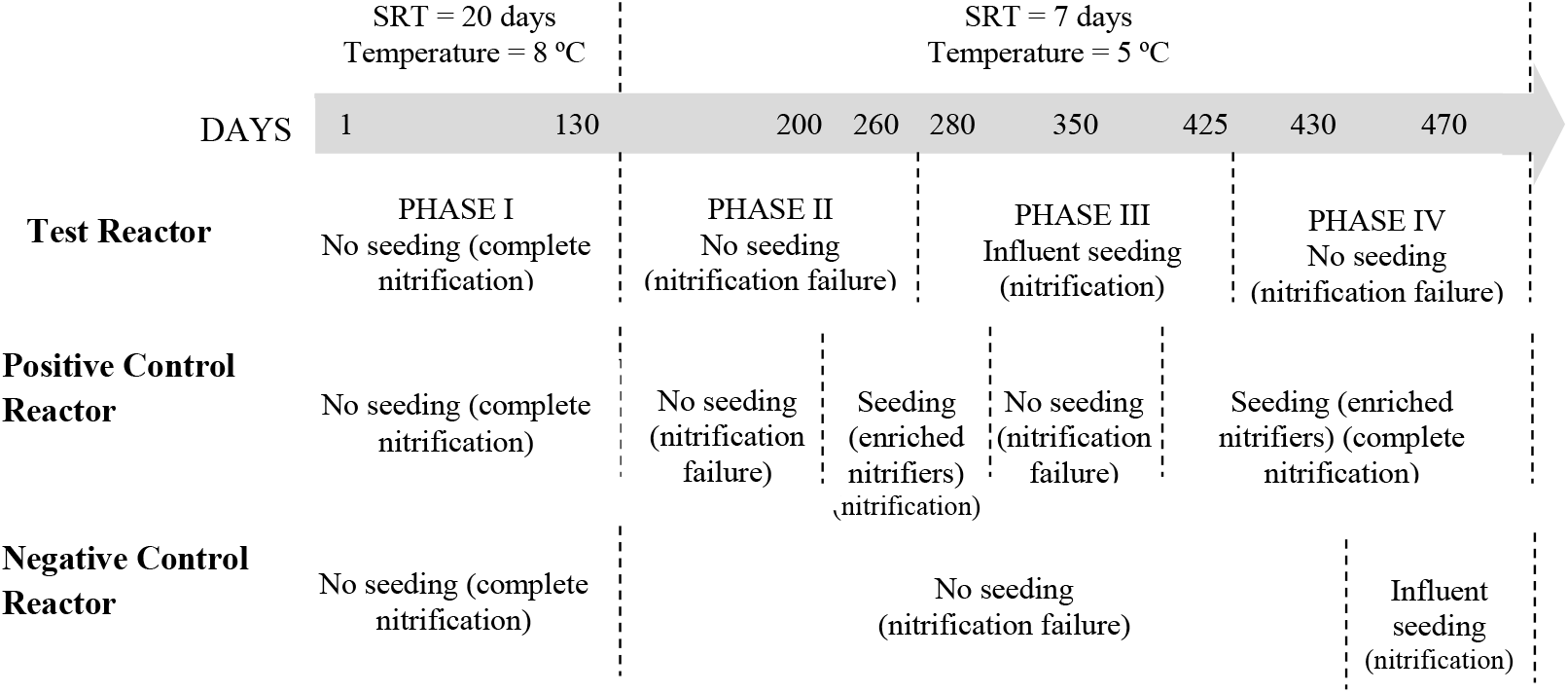
Timeline for seeding of SBRs (from Day 1 to Day 470). The SBRs were operated at 8 °C and at a SRT of 20 days for the initial 130 days. The temperature and SRT were subsequently reduced to 5 °C and 7 days, respectively for the rest of the operation period. Key days are identified on the timeline arrow to indicate start or halt in seeding. Seeding experiments on the Test Reactor were performed in 4 phases as indicated below the timeline. The seeding schedule for the control reactors are also displayed under the timeline bar. Biomass samples were collected from the SBRs on day 40, 70, 150, 200, 230, 270, 312, 354, 400, 430, 450 and 470 for bacterial community analysis. Respective seed biomass samples were also collected in parallel.

The seeding experiments for the Test Reactor were conducted in four phases: (i) Phase I - no seeding (complete nitrification at beginning of the experiment), (ii) Phase II - induction of nitrification failure, (iii) Phase III - seeding of reactor with municipal influent solids, and (iv) Phase IV - reversing the seeding regime between the Test and Negative Control reactors in order to show reproducibility of the influent seeding effect. The timeline for the seeding experiment on the control reactors is also shown in Fig. 2. The nitrification performance was assessed by monitoring the concentration of ammonium (NH_4_^+^) and nitrite (NO_2_^-^)/Nitrate (NO_3_^-^) in the influent and effluent. NH_4_^+^ was determined based on the Berthelot method involving the reaction of NH_4_^+^ as monochloramine with hypochlorite and phenol at a pH of 13 (Rhine et al. 1998). The concentration of NO_2_^-^/NO_3_^-^ was measured based on azo dye formation and reduction of NO_3_^-^ using hydrazine sulphate (Shand et al. 2008). Both tests were performed by microplate assays using the SPECTRAmax^®^ microplate spectrophotometer (CA, USA). The spectrophotometer was operated using the SOFTmax^®^ PRO software. The level of the mixed liquor volatile suspended solids (MLVSS) was monitored according to Standard method 2540 E (APHA 2005).

### 2.3. Bacterial community composition

Thirty-six activated sludge samples were collected across the duration of the study to assess the impact of seeding on the bacterial community composition of the SBRs. Seed samples (influent and cold-adapted nitrifier-enriched biomass) were also collected for analysis. Samples were centrifuged at 10,000×g in eppendorf tubes for 30 minutes using a microcentrifuge (Thermo Scientific, Sorvall Legend Micro 21R, USA) and preserved at −80 °C until time of analysis. A quantity of 0.25 g of wet centrifuged sample was used to extract genomic DNA using the PowerSoil^®^ DNA Isolation Kit (*MO BIO* Laboratories, Inc., Carlsbad, CA). Diluted gDNA (12 ng/mL) was used to amplify the hypervariable V3-V4 region of the *16S rRNA* gene to assess the overall bacterial community composition of the SBRs and their respective seeds. A set of 3 barcoded forward primers targeting the V3 region (*E. coli* position: 338) (Pinto and Raskin 2012) and 1 reverse primer targeting the V4 region (*E. coli* position: 802) (Claesson et al. 2010) were used. Functional genes involved in nitrification were also amplified to identify the nitrifying bacterial profiles of the SBRs and their corresponding seeds. Ammonia oxidizing *bacteria* (AOB) were studied by targeting the *amoA* gene using the barcoded forward primer *amoA-1F* and reverse primer *amoA-2R* (Rotthauwe et al. 1997). Nitrite oxidizing bacteria (NOB) were studied by targeting the *nxrB* gene of the *Nitrospira* genus, considered as the dominant NOB population in activated sludge systems (Hovanec et al. 1998), using the barcoded forward primer *nxrB-F169* and reverse primer *nxrB-638R* (Maixner 2009). The primer sequences and thermal cycles are provided in Table S1 of the Supplementary Data sheet. Each 50μl of PCR reaction mixture contained 0.5 μM of forward and reverse primer each, 1× 5X Bioline PCR colorless buffer (Taunton, MA, USA), 2.75 mM MgCl_2_, 250 μM dNTP (each), 12 ng/mL DNA template and 2.5 units Bioline Taq DNA Polymerase (Taunton, MA, USA) in UltraPure™ DNase/RNase-Free Distilled Water (Invitrogen, Carlsbad, USA).The PCR amplicons were purified using the *MO BIO* UltraClean™ PCR CleanUp Kit (Carlsbad, CA). The quality of the PCR amplicons was assessed using the DNA 1000 microfluidic chip (Agilent Technologies, USA) and the Bioanalyzer 2100 (Agilent Technologies, USA). Any amplicon contaminated with traces of non-specific fragments was further purified using the Agencourt AMPure™ PCR purification kit (Beckman Coulter, Beverly, MA, USA). The concentration of each PCR amplicon was determined using the Quant-iT™ PicoGreen kit (Invitrogen, Carlsbad, USA) and normalized to a concentration of 30 ng/μl. Amplicon mixtures were subjected to emulsion PCR based on Roche-454 Life Science Protocol and pyrosequenced by the GS FLX Titanium Sequencing system using the unidirectional sequencing Lib-L chemistry. Sequencing was performed at the McGill University and Génome Québec Innovation Centre (Montreal, QC).

The *16S rRNA, amoA* and *nxrB* fasta sequence data, flow and quality files were retrieved from raw Standard Flowgram Format files, de-multiplexed, trimmed and filtered using the QIIME software (Caporaso et al. 2010) to retain only good quality sequences devoid of primers and barcodes. USEARCH was used for sequence denoising and chimera removal (both *de novo* and reference-based). Additionally, chimeric sequences were analyzed with UCHIME using default parameters, and were removed. Quality filtered sequences (minimum read length of200 bp, quality score > 25, and without ambiguous bases and mismatches) were clustered at 97% sequence similarity, considered to be related to species level (Islam et al. 2015), into operational taxonomic units (OTUs) using the Uclust algorithm (Edgar 2010). Resulting clusters were subjected to *alpha* and *beta* diversity analyses using default scripts in QIIME and *BiodiversityR* package of the R-software, version 3.2.1. Phylogenetic trees displaying the position of the most abundant AOB and *Nitrospira*-related NOB species were constructed using FastTree (Price et al. 2009) and visualized using FigTree v1.4.2. Taxonomic identification of dominant nitrifying bacterial OTUs was performed using the Functional Gene Pipeline and Repository of the Ribosomal Database Project (Fish et al. 2013), and BLASTN DNA query search (Altschul et al. 1990).

In order to assess the similarity between the influent and mixed liquor nitrifying bacterial profiles, the percentage of sequence reads shared between the two sample matrices was computed using the equation below.

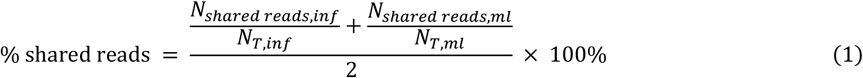

Where *N_shared reads,inf_* is the no. of reads of shared OTUs from the influent, *N_shared reads,ml_* is the no. of shared OTUs from the mixed liquor, *N_T,inf_* is the total no. of reads from the influent and *N_T,ml_* is the total no. of reads from the mixed liquor.

## 3. Results

### 3.1. Impact of influent seeding on nitrification

During the period Day 1-130, all reactors were operated at 8 °C and a SRT of 20 days without seeding, and nitrification of ammonia was almost complete. During Day 131-260, the temperature was decreased to 5 °C, and the SRT was gradually reduced to 7 days. This resulted in the washout of nitrifiers and an increase in effluent NH_4_^+^ concentration to an average of 25 mg/L (Fig. 3) in all reactors. At the same time, the effluent NO_2_^-^/NO_3_^-^ concentration dropped to almost 0 mg-N/L. During Day 261-425, seeding the Test Reactor with approximately 5 mg-COD/L of nitrifying biomass by adding influent solids collected at the LaPrairie full-scale WRRF, decreased the effluent NH_4_^+^ concentration to about 3 mg-N/L and increased NO_2_^-^/NO_3_^-^ concentration to around 20 mg/L, while the Negative Control reactor remained under a state of nitrification failure with high levels of NH_4_^+^ and almost no production of NO_2_^-^/NO_3_^-^ (Fig. 3). This level of influent nitrifier seeding (5 mg-COD/L of nitrifying biomass) was previously determined in the influent of the RAEBL WRRF by respirometry (Jauffur et al. 2014). The constant seeding of the Test Reactor with influent solids maintained nitrification at 5 °C and a SRT of 7 days. During phase IV (Day 426-470), a halt in the supply of influent solids led the effluent NH_4_^+^ concentration to increase rapidly, indicating that nitrifying bacteria were being washed out of the SBR system. Seeding the Negative Control reactor during Day 430-470 with influent solids from the full-scale WRRF effectively caused the effluent NH_4_^+^ level to drop with a corresponding increase in effluent NO_2_^-^/NO_3_^-^ concentration (Fig. 3). These observations provide evidence of the presence of nitrifiers in raw wastewater which supported the nitrifying activity of the reactors.

**Fig. 3.**
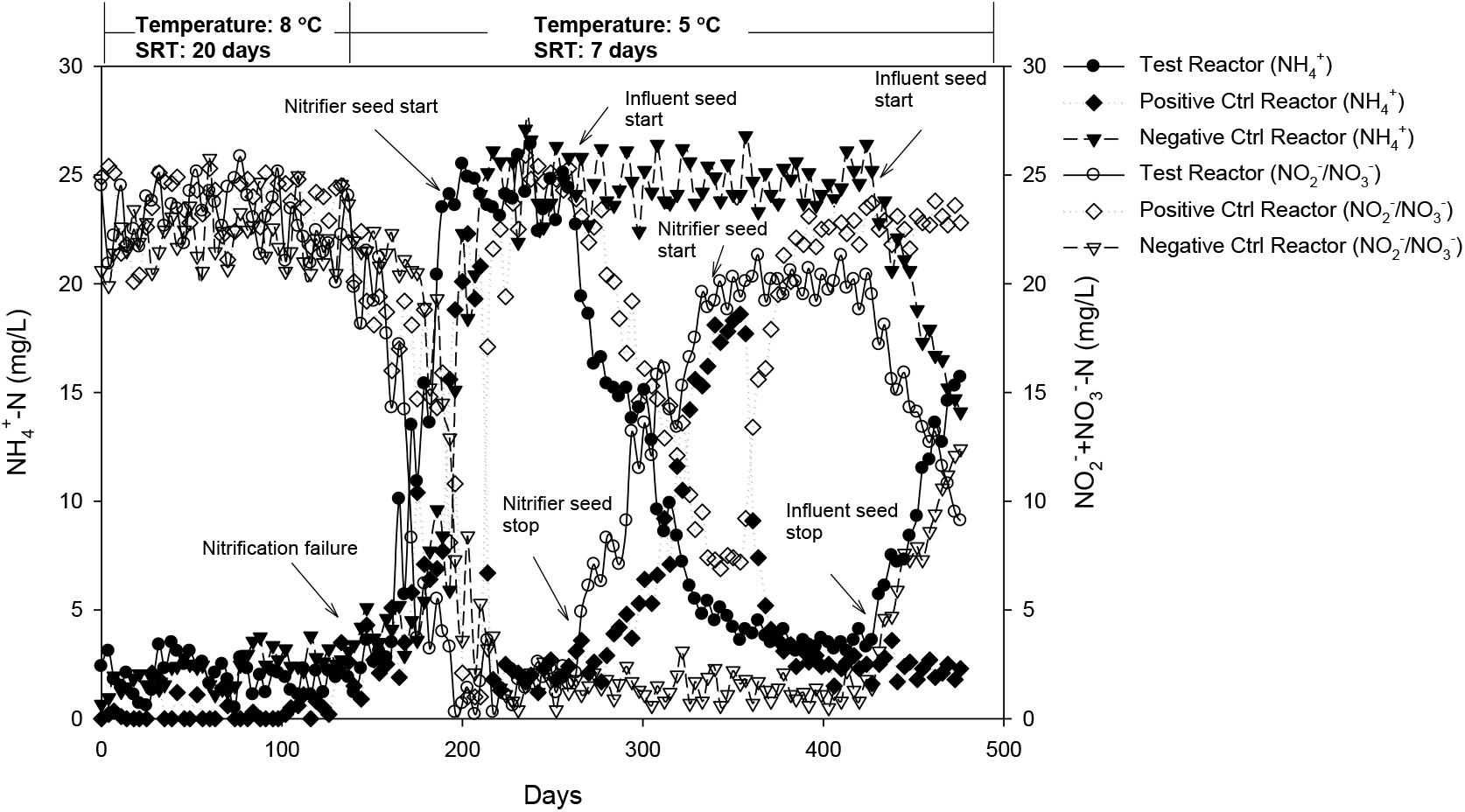
Nitrification performance of SBRs showing effluent NH_4_^+^ and NO_2_^-^/NO_3_^-^ concentrations. Nitrification failure was induced at Day 131 by reducing the operational temperature from 8 °C to 5 °C and SRT from 20 to 7 days. The Test Reactor (Days 261-425) and the Negative Control reactor (Days 430-470) received influent solids as nitrifier seed (Influent seed). The Positive Control reactor (Days 200-279 and Days 350-470) was supplemented with cold-adapted enriched nitrifying biomass as seed (Nitrifier seed). Other operational and performance parameters are provided in Fig. S1 of the Supplementary Data Sheet.

Seeding the Positive Control reactor at Day 200, with the same concentration of approximately 5 mg-COD/L of nitrifying biomass as for the Test Reactor, with cold-adapted nitrifying culture resulted in the re-establishment of near complete nitrification in this reactor. However, the drop in NH_4_^+^ concentration in the effluent occurred within 17 days, while the NH_4_^+^ drop occurred within 75 days for the Test Reactor that received real municipal wastewater solids (Fig. 3). Stopping the supply of nitrifying enrichment at Day 280 led to a gradual increase in effluent NH_4_^+^ concentration, while resuming seeding as from Day 350 restored nitrification. This shows that the supply of external seed was necessary to maintain nitrification in this reactor under unfavorable operational conditions (low temperature and SRT).

### 3.2. Impact of seeding on nitrifying populations

Pyrosequencing of *amoA* and *nxrB* PCR amplicons from reactor and seed samples yielded 121,765 and 121,498 effective sequence tags for AOB and *Nitrospira*-related NOB, respectively. The sequence reads were clustered at 3% cut-off level to generate distinct species-level OTU phylotypes (Keijser et al. 2008). The number of sequence reads, OTU richness and diversity indices for each sample are shown in Table S2. AOB populations showed higher richness as compared to *Nitrospira*-related NOB populations, considering that only the *Nitrospira* genus was studied as NOB. The diversity, as measured by the Hill number (based on Shannon entropy), was fairly low in all reactor samples ranging from 2.08-9.32, and 1.73-4.85 effective number of species for AOB and *Nitrospira*-related NOB populations, respectively. This was due to the extreme dominance of 1-4 OTUs (Fig. 4, S2 and S3).

**Fig. 4.**
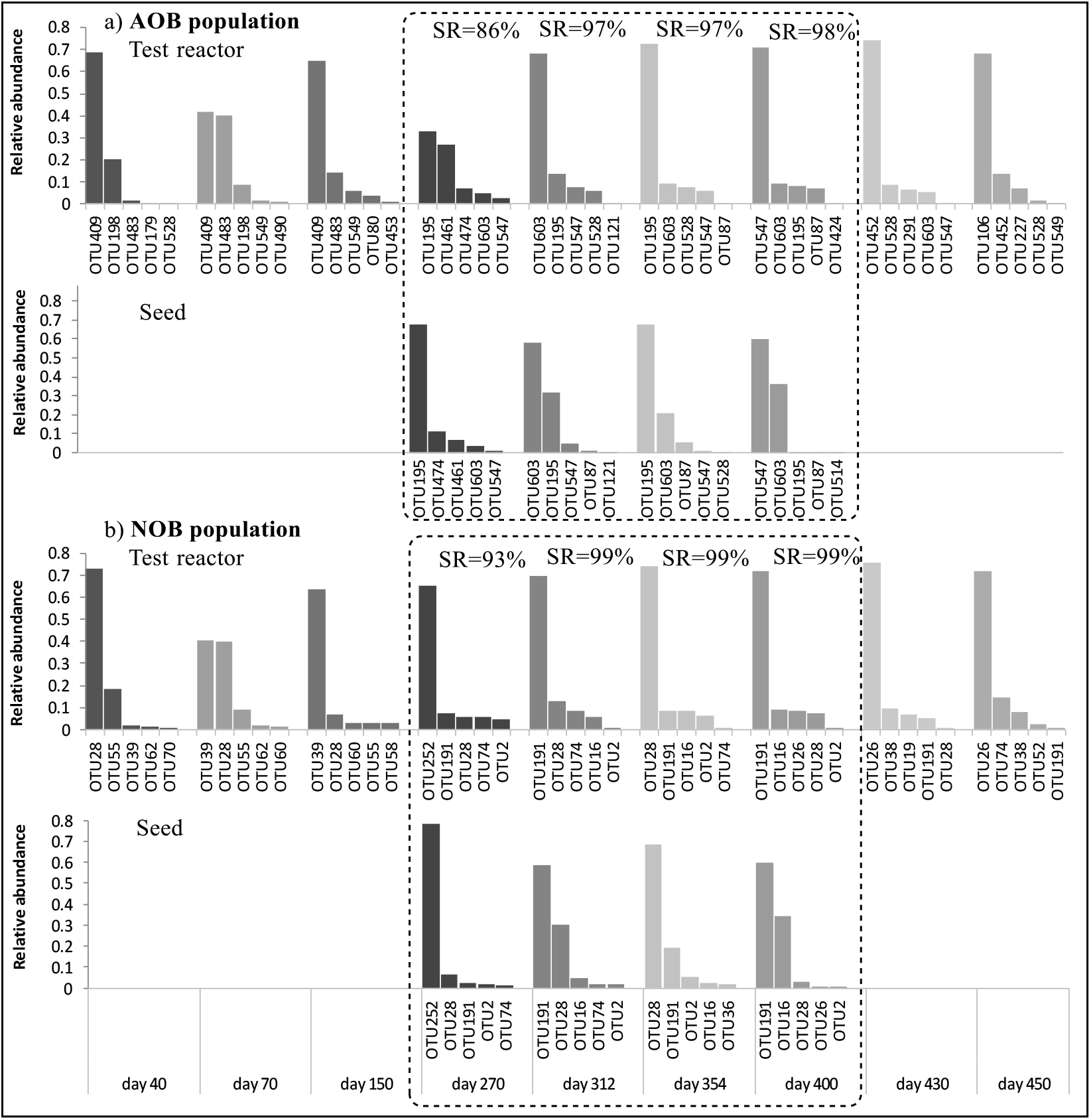
Bar charts showing the 5 most abundant OTUs for AOB (a) and NOB (b) populations in the Test Reactor and its corresponding influent seeds. Seeded activated sludge samples were collected on day 270, 312, 354 and 400, respectively for bacterial community analysis (indicated by the dashed-line box). SR – Shared Reads (Eq.5.1).

During the first experimental period (Days 1-130) when nitrification was sustained within the SBRs because of the 20-d SRT, the most abundant nitrifying populations remained mostly the same for all samples. A certain dynamic was observed in the population structures with the rank-order of OTU abundance, which sometimes varied between samples. When seeding was applied, the nitrifying bacterial population structures of the mixed liquors were very close to the nitrifying population structures of the corresponding seeds (Fig. 4, S2 and S3). This remain true even if the structures of the seed samples varied over time, suggesting that the dynamics of the nitrifying bacterial structure in the mixed liquor were determined by the dynamics of the nitrifying population structure of the seed during the experimental period. Quantitative analyses (based on Eq. 1) showed that 86-98% of the *amoA* sequences were from AOB OTUs shared between the mixed liquor and seed samples collected on the same day. For *Nitrospira*-related NOB, 93-99% of *nxrB* sequences were shared between matched mixed liquor and seed samples. Therefore, seeding had a strong impact in shaping the nitrifying population structure of the mixed liquors for all the reactors.

The taxonomic affiliations of the most abundant AOB and *Nitrospira*-related NOB OTUs were identified (Fig. S4). For the AOB populations, species from five distinct lineages (lineage 1, 2, 3, 5 and 6) could be identified on the phylogenetic tree. Prior to seeding (Day 1-130), *Nitrosomonas oligotropha* (OTU409) was the most abundant AOB species in the Test Reactor. At the time of seeding, *Nitrosomonas europaea* (OTU195) and *Nitrosomonas eutropha* (OTU603) emerged as the most dominant AOB species in the mixed liquor of the Test Reactor. Concurrently, they were also the most abundant AOB species in the influent seed. The same dynamics were observed when seeding the Negative Control reactor at the end of the experimental period. Seeding it with influent solids caused the sewage-derived AOB species, *Nitrosomonas europaea* (OTU 195) and *Nitrosomonas eutropha* (OTU603), to emerge as the most dominant AOB species.

Similar observations were made for *Nitrospira-related* NOB. Prior to seeding, *Ca*. Nitrospira defluvii (OTU28) and uncultured *Nitrospira* sp. (OTU39) were the most dominant *Nitrospira-* related NOB in the Test Reactor. However, seeding the reactor with influent solids caused a shift in the population structure with uncultured *Nitrospira* sp. (OTU252 and OTU191) emerging as the most abundant *Nitrospira*-related NOB. This reflected the *Nitrospira*-NOB population structure of the influent seeds which were also dominated by the same species. Similar observation was made for the Negative Control reactor where the most abundant species namely uncultured *Nitrospira* sp. (OTU39) and *Ca*. Nitrospira defluvii (OTU28) were superseded by the dominant sewage-derived *Nitrospira*-related NOB namely uncultured *Nitrospira* sp. (OTU252 and OTU191). Comparable dynamics were observed with the Positive Control reactor where the same nitrifying AOB species (*Nitrosomonas oligotropha*, OTU409) dominated the mixed liquor during the initial phase of the experiment. Supplementing the reactor with cold-adapted enriched nitrifying biomass caused *Nitrosomonas europaea* (OTU195) and *Nitrosomonas eutropha* (OTU603), to become the most abundant AOB species in the reactor. The seed also induced dominance of sewage-derived *Nitrospira-related* NOB in the mixed liquor of the Positive Control reactor.

### 3.3. Microbial community structures by 16S rRNA gene sequence analysis

Addition of sewage solids to the Test Reactor influenced the heterotrophic bacterial community structures of the mixed liquor. The influent seeding increased the species richness and diversity of the activated sludge communities as shown in Table S3 of the Supplementary Data Sheet. A halt in the supply of influent seed decreased the richness and diversity of the reactor communities. Similarly, the species richness and diversity of the Negative Control reactor increased at the end of the experimental period when influent seed was supplied to the reactor (Table S3).

Principal Coordinate Analysis (PCoA) was used to visualize overall patterns of dispersion of bacterial communities of the seed and reactor samples (Fig. 6). The mixed liquor samples from the three reactors obtained during the first experimental period (D1-D130) clustered in the top-right quadrant, while the influent and nitrifying enrichment seed samples clustered in the bottom- and top-left quadrants, respectively. This indicates that there were clear differences in the composition of the microbial communities. For the Test and Negative Control reactors, the onset of influent seeding induced gradual vectorial drifts of the activated sludge communities towards the seed communities. In the case of the Test Reactor, the drift seemed to be reversible as the community structure moved back towards the initial location on the PCoA plot for the period when supply of influent seed was halted (D430-470). The seed-induced drift was also apparent for the Positive Control reactor which displayed clear gradients following successive addition of enriched nitrifying biomass. When seeding was stopped between Day 312 to 354, the mixed liquor community structure drifted back toward the original cluster but resumed its movement towards the nitrifying enrichment when seeding was restored.

The families which contributed significantly to the ordination have vectors outside the equilibrium circle as shown in Fig. 5. In the bottom-left quadrant, the *Moraxellaceae* and *Campylobacteraceae* families were found particularly imposing on the ordination pattern and predominantly present in influent samples. Lee et al. (2015) also observed their dominance in raw sewage. Opposite to these two vectors were the *Rhodocyclaceae* and *Flavobacteriaceae* families which mapped in the top-right quadrant close to the reactor samples. These families commonly occur in activated sludge (Zhang et al. 2011). The occurrence of *Rhodocyclaceae* and *Flavobacteriaceae* vectors at nearly 180° from the *Moraxellaceae* and *Campylobacteraceae* indicates a negative correlation between them. Seeding of the Test and Negative Control reactors with influent solids decreased the abundance of *Rhodocyclaceae* and to a lesser extent the abundance of *Flavobacteriaceae*, while increasing the abundance of *Moraxellaceae* and *Campylobacteraceae* (Fig. S5). A halt in the supply of influent seed induced the reverse trend by increasing the abundance of *Rhodocyclaceae* and *Flavobacteriaceae* and decreasing the level of *Moraxellaceae* and *Campylobacteraceae* in the mixed liquor. This shows the impact of the influent seeds on structuring the bacterial communities in the activated sludge. Even at the phylum level, this effect was somewhat noticeable where prior to seeding, the mixed liquor of the Test Reactor comprised of *Bacteroidetes* as the predominant phylum consisting of 62.7% of the detected OTUs, followed by *Proteobacteria* (25.6%). Supplementing influent solids, which consisted of predominantly *Proteobacteria* (50.1%) followed by *Bacteroidetes* (18.4%), reduced the abundance of *Bacteroidetes* in the Test Reactor by 15% and increased the level of *Proteobacteria* by 34%. Similar observation was made with the Negative Control reactor where the addition of influent seed reduced the abundance of the *Bacteroidetes* by 25% and increased the level of *Proteobacteria* by 5% in the mixed liquor.

**Fig. 5.**
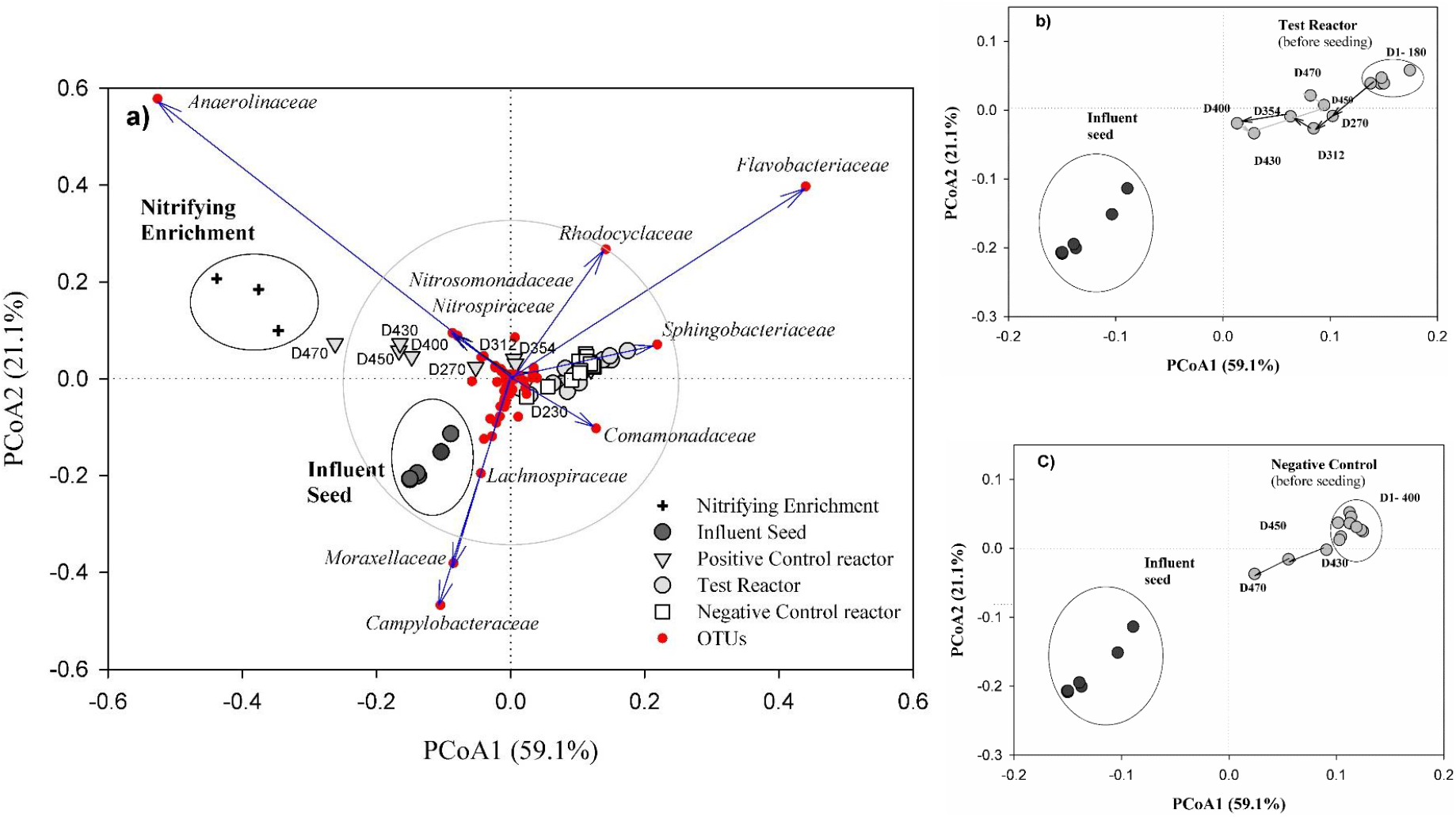
Principal coordinate analysis (PCoA) of *16S rRNA* gene amplicon pyrosequencing data for samples of mixed liquors from Test, Positive Control and Negative Control reactors, and influent and nitrifying enrichment seeds (a). Ordination of samples was based on Bray-Curtis distance at 3% cut-off level. The families to which the identified species belong to are projected on the PCoA plot as red dots. Samples are represented by different symbols. The grey circle corresponds to the equilibrium circle (Legendre and Legendre 1998). Zoomed in portion of PCoA for Test Reactor (b) Negative Control reactor (c) are presented. Numbers starting with “D” indicate the days on which the samples were collected.

The relationship between the influent, sludge of the Test Reactor during seeding, and Negative Control in the absence of seeding during the period Day 270-400 was further explored to determine the number of bacterial genera shared between them. Based on 168 identified genera, 48 were shared among the influent, Test Reactor and Negative Control reactor (Fig. S6). Although the influent had distinct microbial community composition, it imparted 22% of the number of genera to the Test Reactor including *Nitrosomonas* and *Nitrospira* which were key members in reestablishing nitrification during the failure period. This shows that despite their marked differences, bacterial communities of the activated sludge reactor were not totally independent from influent wastewater communities.

## 4. Discussion

### 4.1. Influent-induced nitrification

Addition of influent solids to the Test Reactor operated at near washout conditions (temperature of 5 °C and SRT of 7 days) restored nitrification causing a decrease in effluent NH_4_^+^ level and a concurrent increase in effluent NO_2_^-^/NO_3_^-^ concentration. Interrupting the supply of influent seed to the Test Reactor caused the effluent NH_4_^+^ level to increase and the effluent NO_2_^-^/NO_3_^-^ level to decrease indicating that a constant supply of influent seed was necessary to maintain nitrification in the Test Reactor. Reversing the seeding regime by supplementing the Negative Control reactor with influent seed restored nitrification showing that the influent seeding effect was reproducible, and a certain level of operational control could be exerted on the process.

This provides strong evidence that incoming raw wastewater contains nitrifying bacteria. It also demonstrates that the phenomenon of natural nitrifier seeding is playing a key role in sustaining nitrification at full-scale level in extreme conditions such as low temperature and SRT. Since the metabolic induction time of influent nitrifiers have been shown to be only a few hours (Jauffur et al. 2014), and considering that wastewater treatment systems are normally operated at a few day SRT, the seeded nitrifiers can be adsorbed on activated sludge flocs to bioaugment nitrifying populations. Such adsorption is likely since nitrifiers have been shown to possess strong adhesion properties. Larsen et al. (2008) studied the adsorption of *N. oligotropha* and *Nitrospira* sp. and found that activated sludge flocs consisted of an easily detachable fraction (5-15%), a fraction resistant to deflocculation (15-40%), and a strong, non-detachable fraction (50-75%). Nitrifiers presumably belong to the non-detachable fraction of activated sludge flocs.

Supplementing influent seed was found to be necessary in order to sustain NH_4_^+^ removal under conditions which were not conducive to nitrification. Such influent-induced regain in nitrification activity may be important in modeling nitrifying activated systems and points to an important limitation of ASM models which currently assume no autotrophic nitrifying biomass in influent wastewater. Integrating the natural seeding of nitrifiers by influent wastewaters may render nitrification model predictions more accurate especially for systems operating under unfavorable conditions such as cold climate or at sub-optimal SRT. In design of nitrifying activated sludge systems, a safety factor of 2-3 is usually applied to account for fluctuations in nitrifiers’ growth rates resulting from specific local conditions (load variation, temperature, pH, inhibition and dissolved oxygen) (Rittmann and McCarty 2001). The supply of a natural nitrifier seed by the influent flow can increase this safety factor without increasing reactor footprint considering that any increase in SRT will increase the solids inventory, which must be offset by increasing the size of the bioreactor and clarifier. In addition, supply of influent seed can help a nitrifying system on the verge of washout to recover by re-establishing its nitrifying population.

Bacterial community analyses showed that the diversity of the bench-scale reactors was lower than the diversity observed at full-scale systems. For instance, the Hill diversity number for AOB populations in the seeded Test Reactor ranged between 3.18-3.71 as compared to full-scale studies where the observed AOB diversity ranged between 6.69-10.07 (Baek et al. 2010) and 13.0-66.0 (Jauffur et al. 2018, Unpublished data). This is due to the presence of only a few dominant species (1-4) in the samples analyzed followed by long tails of rare taxa as shown by the OTU rankabundance analyses. Moreover, discrepancies in bacterial diversities at lab- and full-scale systems have also been reported in other studies and may result from differences in operational and environmental factors (Briones and Raskin 2003).

Most importantly, the patterns of nitrifying bacterial population in the activated sludge of the Test Reactor, and the Negative Control reactor at the end of the experimental period, were similar to the ones of the corresponding influent seeds. The most abundant AOB and *Nitrospira*-related NOB OTUs observed in the influent seeds also occurred as the dominant ones in the activated sludge. This shows that seeding the Test Reactor or Negative Control reactor with influent solids restructured the community of the reactors with *Nitrosomonas europaea* and *Nitrosomonas eutropha* emerging as dominant species in the activated sludge. Previous studies have reported the wide dominance of these species in full-scale WRRFs receiving municipal wastewaters (Figuerola and Erijman 2010, Zhang et al. 2011). Stopping the supply of influent seed to the Test Reactor incurred a restructuring of the most dominant AOB species, a period coinciding with a failure in nitrification. According to (Wittebolle et al. 2005), nitrification failures at full-scale WRRFs are intimately linked to shifts in microbial community structures. For NOB populations, mostly uncultured *Nitrospira* species were found dominant in the activated sludge and influent seeds. Abundant identified species included *Ca*. Nitrospira defluvii and *Nitrospira moscoviensis*, which are characteristic NOB species inhabiting activated sludge (Koch et al. 2015, Kruse et al. 2013). In this case as well, the dominant *Nitrospira*-related NOB species identified in the activated sludge were found to also occur as dominant species in the influent seed. The same dynamics occurred in the Positive Control reactor where addition of cold-adapted enriched nitrifying biomass restructured the nitrifying communities such that the most abundant species in the seed became dominant in the mixed liquor of the reactor.

Based on the above observations, raw wastewater is found to be a key factor shaping nitrifying communities in full-scale wastewater treatment systems. The source of nitrifying bacteria in the influent wastewaters was not investigated in this study. However, it is believed that they enter sewer networks bound to soil particles through surface runoffs. Once in the sewer, they may colonize strategic places conducive for them to grow. Erosion and sloughing of sewer biofilms are potential means of biomass transportation to full-scale WRRFs (Jahn and Nielsen 1998). Considering that influent streams and full-scale bioreactors are physically connected systems, it is not surprising that bacterial species, including nitrifiers, are continuously supplied to aeration basins by influent wastewaters. In this case, we show that the influent is a key source for maintaining activity and community structure of nitrifiers in activated sludge.

### 4.2. Seeding effect on heterotrophic bacterial populations

Unlike nitrifying bacterial populations, heterotrophic communities have a large breadth of ecological functions (Quince et al. 2008). According to Saunders et al. (2015), these different functional entities in activated sludge ecosystems can mainly be classified as abundant core or transient populations. In this context, it would be legitimate to question the role of immigrant microorganisms conveyed by influent wastewaters and assess their impacts on the selection of bacterial populations or on the niche occupation of bacterial communities in activated sludge systems.

The cluster analysis showed that activated sludge and influent seeds had distinct bacterial populations and mapped apart on the PCoA bi-plot. This difference between influent and sludge community compositions has been reported in earlier studies (Hashimoto et al. 2014, Lee et al. 2015, Liu et al. 2007). The influent communities comprised of families such as *Moraxellaceae, Campylobacteraceae* and *Lachnospiraceae* which are typically present in untreated wastewaters as also observed in previous studies (McLellan et al. 2010, Ye and Zhang 2013). The inability of certain influent bacteria to grow and persist in environmental conditions prevalent in activated sludge may explain this dissimilarity. Activated sludge microorganisms have broad carbon-utilization profiles with an enriched community of degraders acclimated to specific bioreactor environments, which would impose a selective growth of specific populations. Based on our observations, it is clear that the bioreactors imposed a selective pressure on the incoming immigrant heterotrophic populations leading to distinct populations as shown on the PCoA plot. Nonetheless, the immigrant communities also appear to modulate the structure of the activated sludge communities. For instance, seeding the Test Reactor with influent solids decreased the abundance of *Rhodocyclaceae* and *Flavobacteriaceae* while increasing the abundance of *Moraxellaceae* and *Campylobacteraceae*. Such negative correlation between these families was also observed when seeding the Negative Control reactor with influent solids. This effect can be visualized on the ordination plot where seeding of the reactors with influent solids induced vectorial drifts of the communities towards their respective seeds but regressed to their initial positions when seeding was halted. This shows that influent microbiome has an impact on shaping activated sludge communities, and argues against Hashimoto et al. (2014) who opined that influent communities have no influence on reactor communities. About 22% of the genera identified in the seeded Test Reactor appeared to come from the influent biomass indicative that raw sewage contributed in structuring the microbial community of the reactor. According to Verberk (2012), these overlapping taxa may be generalists capable of surviving in both sewer and reactor environments. In subsequence, activated sludge communities are not completely independent from influent communities as recognized by Lee et al. (2015).

At full-scale level, raw sewage communities may, thus, be contributing to determine bacterial community compositions and structures of activated sludge systems. Curtis and Craine (1998) found a large number of microorganisms from sewage in mixed liquor samples even though according to them some may not have major functional significance in the treatment process. The supply of microorganisms to full-scale activated sludge systems may be constantly occurring through immigration as observed by Leibold et al. (2004). Our findings provide evidence that influent wastewater influences activated sludge communities by supplying consortia of bacteria, which may be contributing to the treatment of the incoming wastewater.

## 5. Conclusions

Sequencing batch reactors fed with sterile synthetic wastewater (“*Syntho*”) were operated above washout conditions (low temperature and SRT) to intentionally induce nitrification failure. Addition of real influent solids harvested from the full-scale wastewater treatment facility restored nitrification indicating the presence of live nitrifiers in raw sewage. The wastewater solids stabilized the treatment systems and maintained nitrification in an otherwise non-conducive condition. Phylogenetic analyses revealed that the taxonomic identity of the most abundant nitrifiers in the influent wastewater occurred as the most abundant nitrifiers in the corresponding activated sludge. These findings demonstrate the existence of natural seeding of nitrifiers at fullscale wastewater treatment level by incoming raw sewage. The process was successfully replicated in the laboratory where addition of the influent solids restored nitrification in the bench-top reactors operated at washout conditions. It is believed that these nitrifying microorganisms originate primarily from soils and enter combined sewers through surface runoffs adhered to soil particles. Once inside the sewers, the nitrifying populations may be structured based on selective processes impacting on them. Since heterotrophic bacterial populations are also conveyed by raw sewage to full-scale wastewater treatment facilities, it was legitimate to question the role of immigrant heterotrophic populations in structuring the local communities in the bioreactors. Heterotrophic bacterial community structures were distinct in influents and activated sludge mixed liquors. Based on our analyses it was evident that the bioreactors were imposing a selective pressure on the incoming heterotrophs from the real wastewaters, leading to distinct communities. Nonetheless, seeding the reactors with wastewater solids modified the mixed liquor communities, making their structures more similar to the seed. The findings that nitrifiers in raw wastewater act as potential seeds have direct impacts on the conception, design and operation of activated sludge wastewater treatment plants, including the sizing of the bioreactors, considering that nitrification highly influences the determination of the size and energy requirement of activated sludge wastewater treatment systems performing nitrification. Investigating the effect of influent nitrifier seeding on modeling practices may shed light on the accuracy of current parameters and associated uncertainties involved in developing nitrification models.

## Acknowledgements

We thank the plant operators of the RAEBL water resource recovery facility (WRRF), who assisted us in collecting raw sewage and mixed liquor samples. We would also like to thank the reviewers for their review and constructive comments on this manuscript.

## Appendix A. Supplementary data

## SUPPLEMENTARY DATA

**Table S1.**
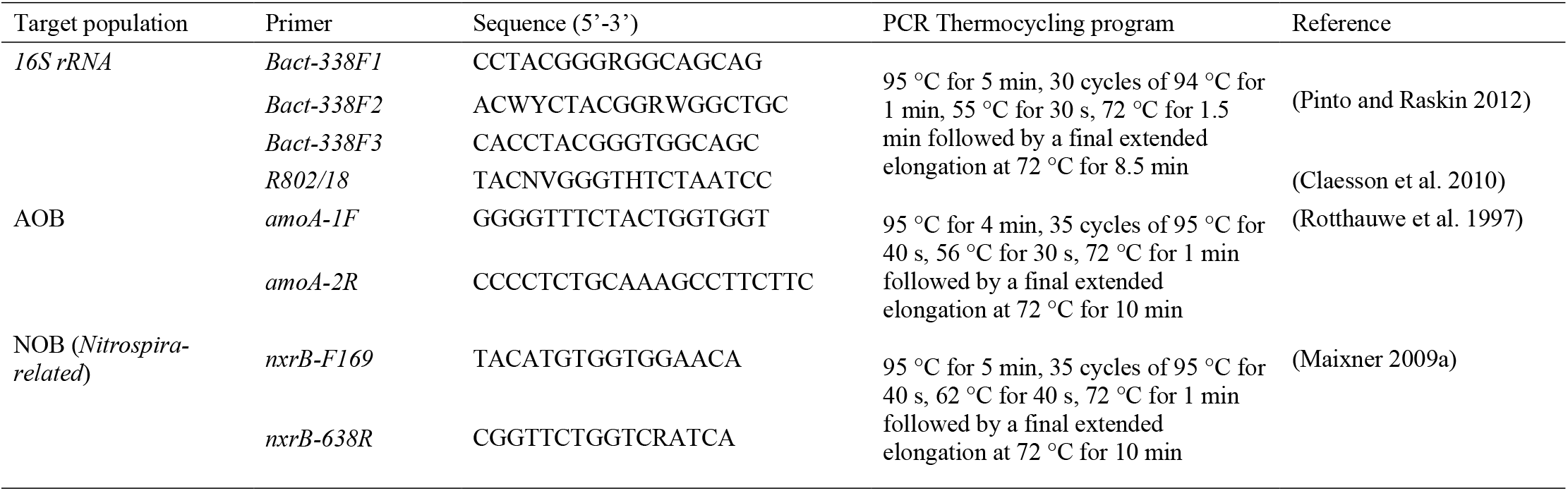
Primers and PCR thermocycling programs used for amplification of *16S rRNA, amoA* and *nxrB* genes.

**Table S2.** Number of reads, OTU richness and diversity indices for autotrophic nitrifying bacterial populations in enriched nitrifier seed, municipal influent and SBR mixed liquor samples determined by *amoA* and *nxrB* PCR amplicon pyrosequencing.

| Samples | Ammonia Oxidizing Bacteria (AOB) using *amoA* |  |  |  |  |  | Nitrite Oxidizing Bacteria (NOB) using *nxrB* |  |  |  |  |  |
| --- | --- | --- | --- | --- | --- | --- | --- | --- | --- | --- | --- | --- |
|  | No. of reads<br>(denoised) | OTU<br>richness | Shannon diversity |  |  | Simpson<br>diversity no. | No. of reads<br>(denoised) | OTU<br>richness | Shannon diversity |  |  | Simpson<br>diversity no. |
|  |  |  | Entropy<br>(nat) | Hill no. | Equitability |  |  |  | Entropy<br>(nat) | Hill no. | Equitability |  |
| Enriched nitrifier seed |  |  |  |  |  |  |  |  |  |  |  |  |
| Day 70 | 3915 | 98 | 1.36 | 3.90 | 0.30 | 2.40 | 3837 | 29 | 1.24 | 3.46 | 0.37 | 2.35 |
| Day 270 | 3321 | 107 | 1.51 | 4.53 | 0.32 | 2.13 | 3001 | 35 | 1.35 | 3.86 | 0.38 | 2.55 |
| Day 400 | 3363 | 98 | 1.01 | 2.75 | 0.22 | 2.11 | 3259 | 33 | 0.97 | 2.63 | 0.28 | 2.13 |
| Influent seed |  |  |  |  |  |  |  |  |  |  |  |  |
| Day 270 | 3033 | 86 | 1.48 | 4.40 | 0.33 | 2.11 | 3466 | 35 | 1.17 | 3.21 | 0.33 | 0.93 |
| Day 312 | 3465 | 119 | 1.19 | 3.28 | 0.25 | 2.27 | 3354 | 33 | 1.12 | 3.07 | 0.32 | 2.27 |
| Day 354 | 3423 | 116 | 1.18 | 3.24 | 0.25 | 1.99 | 3312 | 32 | 1.07 | 2.92 | 0.31 | 1.96 |
| Day 400 | 3347 | 98 | 0.97 | 2.63 | 0.21 | 2.05 | 3270 | 33 | 0.97 | 2.63 | 0.28 | 2.09 |
| Day 430 | 3745 | 123 | 1.33 | 3.79 | 0.28 | 2.42 | 3550 | 42 | 1.26 | 3.54 | 0.34 | 2.58 |
| Day 450 | 3076 | 90 | 1.00 | 2.72 | 0.22 | 2.16 | 3000 | 33 | 1.00 | 2.72 | 0.29 | 2.23 |
| Day 470 | 3441 | 101 | 1.09 | 2.99 | 0.23 | 2.24 | 3323 | 39 | 1.05 | 2.87 | 0.29 | 2.32 |
| Test reactor |  |  |  |  |  |  |  |  |  |  |  |  |
| Day 40 | 3231 | 104 | 1.28 | 3.60 | 0.28 | 1.96 | 4089 | 34 | 0.99 | 2.69 | 0.28 | 1.76 |
| Day 70 | 3130 | 86 | 1.51 | 4.54 | 0.34 | 2.91 | 3045 | 30 | 1.46 | 4.29 | 0.43 | 3.01 |
| Day 150 | 3974 | 79 | 1.48 | 4.39 | 0.35 | 2.25 | 3347 | 32 | 1.35 | 3.86 | 0.39 | 2.42 |
| Day 270 | 3409 | 95 | 1.31 | 3.71 | 0.58 | 5.14 | 3011 | 74 | 1.51 | 4.51 | 0.35 | 2.26 |
| Day 312 | 3083 | 114 | 1.24 | 3.46 | 0.27 | 2.03 | 3965 | 32 | 1.08 | 2.94 | 0.31 | 1.95 |
| Day 354 | 3823 | 114 | 1.16 | 3.18 | 0.24 | 1.83 | 3695 | 36 | 1.25 | 3.49 | 0.35 | 2.15 |
| Day 400 | 3011 | 102 | 1.25 | 3.49 | 0.25 | 1.90 | 3897 | 32 | 1.04 | 2.82 | 0.30 | 1.86 |
| Day 430 | 3237 | 81 | 0.91 | 2.48 | 0.56 | 1.02 | 3128 | 26 | 0.81 | 2.24 | 0.24 | 1.67 |
| Day 450 | 3422 | 62 | 0.73 | 2.08 | 0.50 | 0.81 | 3426 | 19 | 0.55 | 1.73 | 0.18 | 0.78 |
| Positive Control |  |  |  |  |  |  |  |  |  |  |  |  |
| Day 40 | 3521 | 93 | 1.76 | 5.82 | 0.39 | 2.98 | 3413 | 31 | 1.49 | 4.43 | 0.43 | 2.75 |
| Day 70 | 3142 | 83 | 1.55 | 4.71 | 0.35 | 2.82 | 3077 | 36 | 1.44 | 4.22 | 0.40 | 2.31 |
| Day 150 | 3137 | 104 | 2.23 | 9.32 | 0.48 | 3.29 | 3612 | 35 | 1.53 | 4.64 | 0.43 | 2.18 |
| Day 230 | 3356 | 102 | 1.86 | 6.42 | 0.25 | 3.17 | 3809 | 43 | 1.58 | 4.85 | 0.42 | 2.93 |
| Day 270 | 3296 | 118 | 1.92 | 6.82 | 0.40 | 3.24 | 3151 | 36 | 1.49 | 4.44 | 0.42 | 2.32 |
| Day 312 | 3871 | 70 | 1.14 | 3.12 | 0.26 | 1.57 | 4313 | 28 | 1.08 | 2.94 | 0.32 | 1.51 |
| Day 400 | 3847 | 98 | 1.78 | 5.92 | 0.39 | 1.95 | 3756 | 40 | 1.42 | 4.14 | 0.38 | 2.23 |
| Day 430 | 3400 | 107 | 1.64 | 5.16 | 0.35 | 1.93 | 3302 | 46 | 1.52 | 4.57 | 0.40 | 2.67 |
| Day 450 | 3473 | 118 | 1.76 | 5.81 | 0.37 | 2.06 | 3354 | 41 | 1.49 | 4.44 | 0.40 | 2.51 |
| Day 470 | 3802 | 95 | 2.03 | 7.61 | 0.45 | 2.02 | 3707 | 48 | 1.55 | 4.71 | 0.41 | 2.85 |
| Negative Control |  |  |  |  |  |  |  |  |  |  |  |  |
| Day 40 | 3738 | 76 | 1.50 | 4.46 | 0.35 | 2.93 | 3637 | 22 | 1.38 | 3.98 | 0.45 | 2.94 |
| Day 70 | 4467 | 96 | 1.53 | 4.64 | 0.34 | 2.88 | 3380 | 28 | 1.42 | 4.14 | 0.43 | 2.89 |
| Day 150 | 3465 | 74 | 1.39 | 4.03 | 0.32 | 2.01 | 3523 | 33 | 1.22 | 3.38 | 0.35 | 1.95 |
| Day 430 | 3820 | 70 | 0.98 | 2.66 | 0.23 | 1.73 | 3718 | 58 | 0.93 | 2.53 | 0.23 | 1.68 |
| Day 450 | 3820 | 70 | 1.13 | 3.10 | 0.23 | 1.75 | 3754 | 20 | 1.02 | 2.45 | 0.30 | 1.70 |
| Day 470 | 3161 | 80 | 1.25 | 3.50 | 0.25 | 2.05 | 3018 | 25 | 1.10 | 2.56 | 0.29 | 1.94 |

**Table S3.** Number of reads, OTU richness and diversity indices for the entire microbial communities from enriched nitrifier seed, wastewater influent and SBR mixed liquor as determined by *16S rRNA* gene amplicon pyrosequencing.

| Samples | No. of reads (denoised) | OTU richness | Shannon diversity |  |  | Simpson diversity no.<sup>b</sup> |
| --- | --- | --- | --- | --- | --- | --- |
|  |  |  | Entropy (nat) | Hill no.<sup>a</sup> | Equitability |  |
| **Enriched nitrifier seed** |  |  |  |  |  |  |
| Day 70 | 6480 | 114 | 1.84 | 6.28 | 0.39 | 2.43 |
| Day 270 | 5544 | 107 | 2.56 | 12.9 | 0.55 | 5.02 |
| Day 400 | 6246 | 131 | 2.31 | 10.05 | 0.47 | 3.14 |
| **Influent seed** |  |  |  |  |  |  |
| Day 270 | 5599 | 242 | 3.69 | 40.05 | 0.67 | 18.09 |
| Day 312 | 5731 | 237 | 3.71 | 40.89 | 0.68 | 18.60 |
| Day 354 | 6566 | 217 | 3.74 | 42.25 | 0.69 | 19.39 |
| Day 400 | 5930 | 319 | 4.00 | 54.82 | 0.69 | 24.11 |
| Day 430 | 4040 | 205 | 3.69 | 39.86 | 0.69 | 17.82 |
| Day 450 | 5761 | 213 | 3.76 | 42.81 | 0.70 | 20.06 |
| Day 470 | 5478 | 225 | 3.70 | 40.49 | 0.68 | 18.24 |
| **Test reactor** |  |  |  |  |  |  |
| Day 40 | 5583 | 143 | 2.66 | 14.32 | 0.54 | 6.48 |
| Day 70 | 5943 | 151 | 2.59 | 13.28 | 0.52 | 6.98 |
| Day 150 | 5530 | 165 | 3.03 | 20.73 | 0.59 | 9.51 |
| Day 200 | 5010 | 177 | 2.95 | 19.19 | 0.57 | 8.18 |
| Day 230 | 5530 | 174 | 3.18 | 23.98 | 0.62 | 9.41 |
| Day 270 | 5321 | 214 | 3.36 | 28.79 | 0.63 | 11.38 |
| Day 312 | 5942 | 227 | 3.42 | 30.57 | 0.65 | 12.45 |
| Day 354 | 6611 | 225 | 3.39 | 29.55 | 0.64 | 12.39 |
| Day 400 | 6826 | 210 | 3.35 | 28.50 | 0.63 | 11.52 |
| Day 430 | 6843 | 189 | 3.04 | 20.91 | 0.58 | 8.31 |
| Day 450 | 6671 | 164 | 3.02 | 20.49 | 0.59 | 9.52 |
| Day 470 | 6033 | 160 | 2.96 | 19.25 | 0.58 | 8.26 |
| **Positive Control** |  |  |  |  |  |  |
| Day 40 | 5788 | 151 | 3.08 | 21.83 | 0.61 | 9.80 |
| Day 70 | 5016 | 139 | 2.71 | 14.98 | 0.55 | 7.05 |
| Day 150 | 5652 | 143 | 2.81 | 16.62 | 0.57 | 6.32 |
| Day 200 | 6579 | 178 | 3.39 | 29.66 | 0.65 | 10.58 |
| Day 230 | 5536 | 204 | 3.56 | 35.10 | 0.67 | 15.14 |
| Day 270 | 5945 | 210 | 3.62 | 37.34 | 0.68 | 15.23 |
| Day 312 | 6794 | 170 | 3.07 | 21.58 | 0.59 | 8.65 |
| Day 354 | 5454 | 144 | 2.54 | 12.68 | 0.51 | 4.75 |
| Day 400 | 5871 | 208 | 3.62 | 37.34 | 0.68 | 15.35 |
| Day 430 | 5245 | 211 | 3.71 | 40.85 | 0.69 | 15.42 |
| Day 450 | 5303 | 214 | 3.85 | 46.99 | 0.72 | 15.73 |
| Day 470 | 5880 | 217 | 3.88 | 48.42 | 0.69 | 15.81 |
| **Negative Control** |  |  |  |  |  |  |
| Day 40 | 5541 | 95 | 2.73 | 15.40 | 0.60 | 8.85 |
| Day 70 | 5448 | 125 | 2.69 | 14.88 | 0.56 | 8.32 |
| Day 150 | 5677 | 127 | 2.62 | 13.79 | 0.54 | 7.19 |
| Day 200 | 5010 | 128 | 2.98 | 19.62 | 0.61 | 10.42 |
| Day 230 | 5309 | 133 | 2.94 | 18.87 | 0.60 | 10.12 |
| Day 270 | 5900 | 129 | 2.88 | 17.74 | 0.59 | 8.02 |
| Day 312 | 5968 | 151 | 3.12 | 22.73 | 0.62 | 10.84 |
| Day 354 | 7024 | 148 | 3.27 | 26.22 | 0.65 | 13.81 |
| Day 400 | 5097 | 144 | 3.27 | 26.24 | 0.66 | 14.72 |
| Day 430 | 5076 | 239 | 3.58 | 35.89 | 0.65 | 16.18 |
| Day 450 | 5075 | 231 | 3.74 | 42.06 | 0.69 | 19.37 |
| Day 470 | 5182 | 241 | 3.49 | 32.96 | 0.64 | 19.42 |

<sup>a</sup> Hill no. = exp [Shannon entropy]
<sup>b</sup> Simpson diversity no. = $1 / \sum_{i=1}^S p_i^2$ , where $p_i$ is the proportion of the $i^{th}$ OTU

**Fig. S1.**
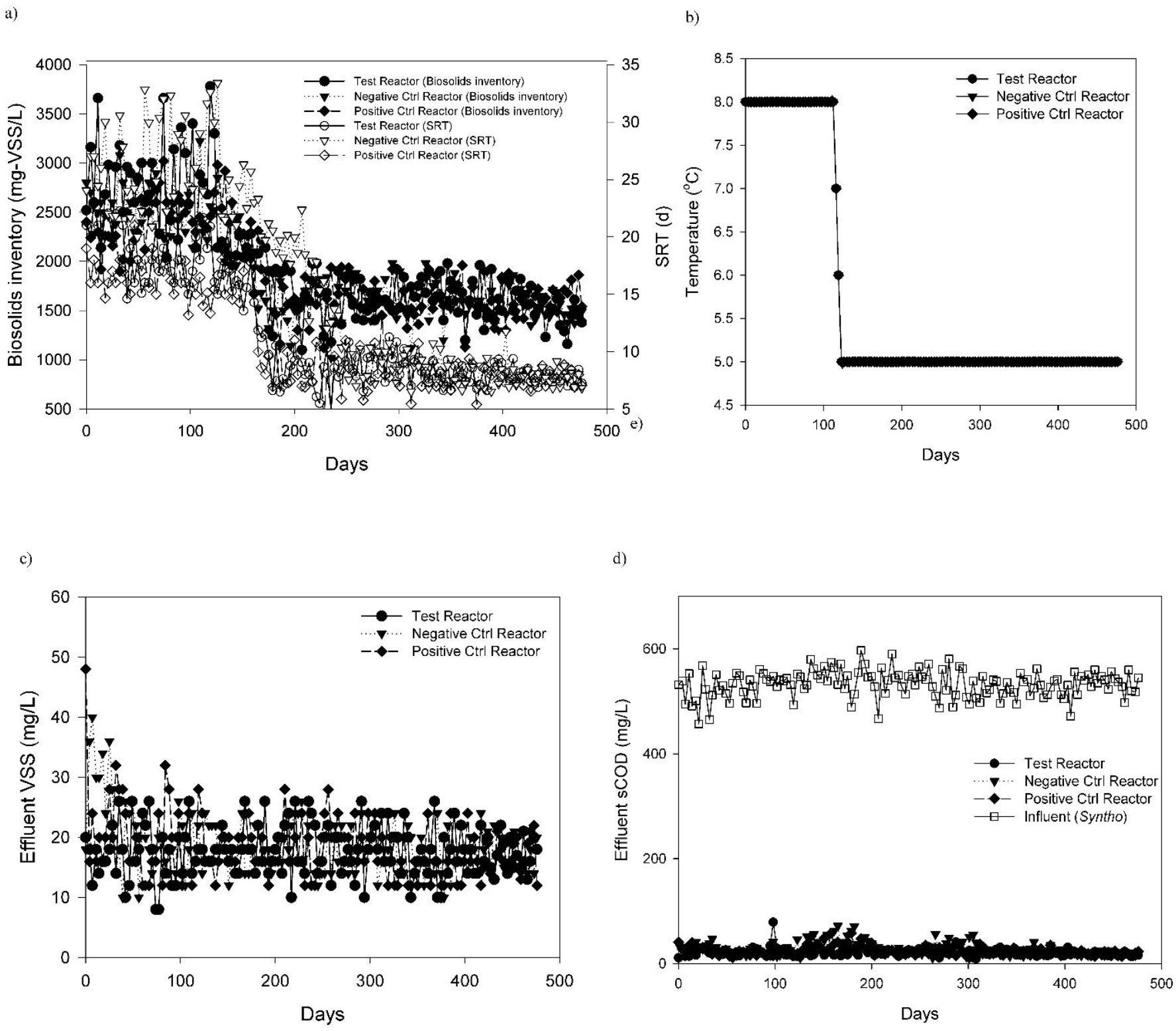
Operational and performance parameters for SBRs. Variation of biosolids inventory and SRT (a), temperature (b), effluent VSS (c), and effluent sCOD (d) over time.

**Fig. S2.**
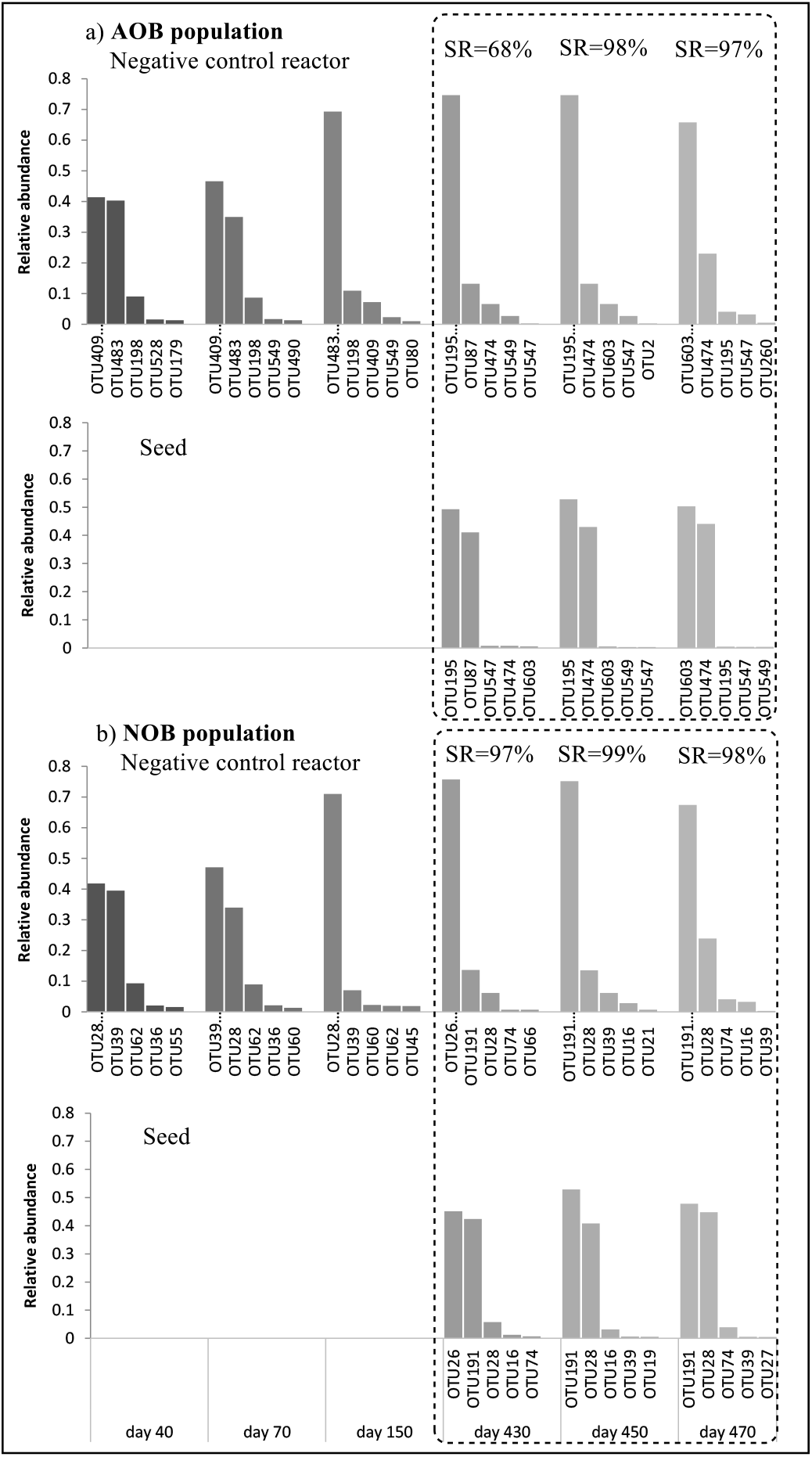
Bar charts showing the 5 most abundant OTUs for AOB (a) and NOB (b) populations in the Negative Control reactor and its corresponding seed. Seeding of the reactor with influent biomass was performed on day 430, 450 and 470, respectively (indicated by the dashed-line box). SR - Shared Reads (Eq. 1).

**Fig. S3.**
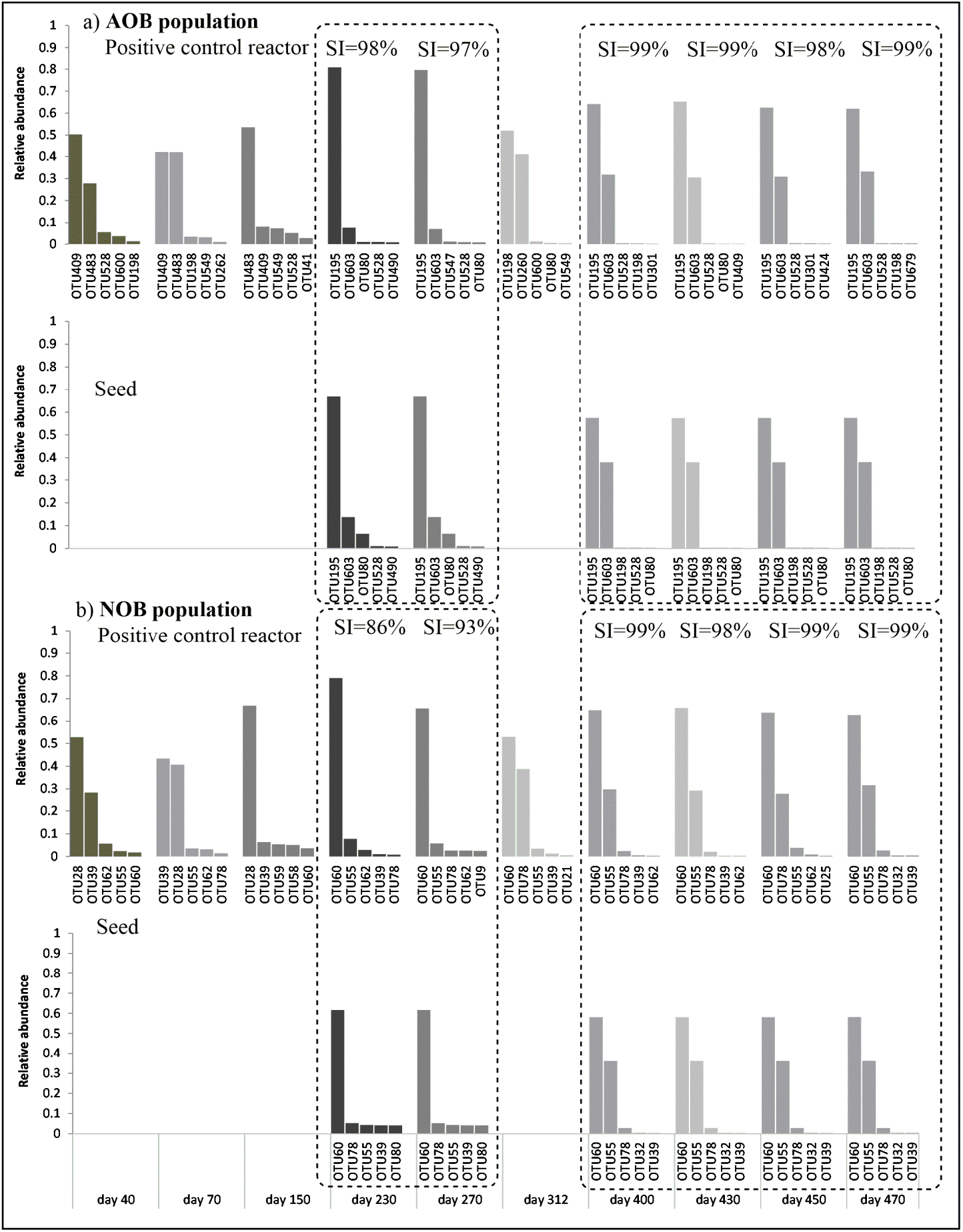
Bar charts showing the 5 most abundant OTUs for AOB (a) and NOB (b) populations in the Positive Control reactor and its corresponding seed. Seeding of the reactor with enriched cold-adapted nitrifying biomass was performed on day 230, 270, 400, 430, 450 and 470, respectively (indicated by the dashed-line box). SR - Shared Reads (Eq. 1).

**Fig. S4.**
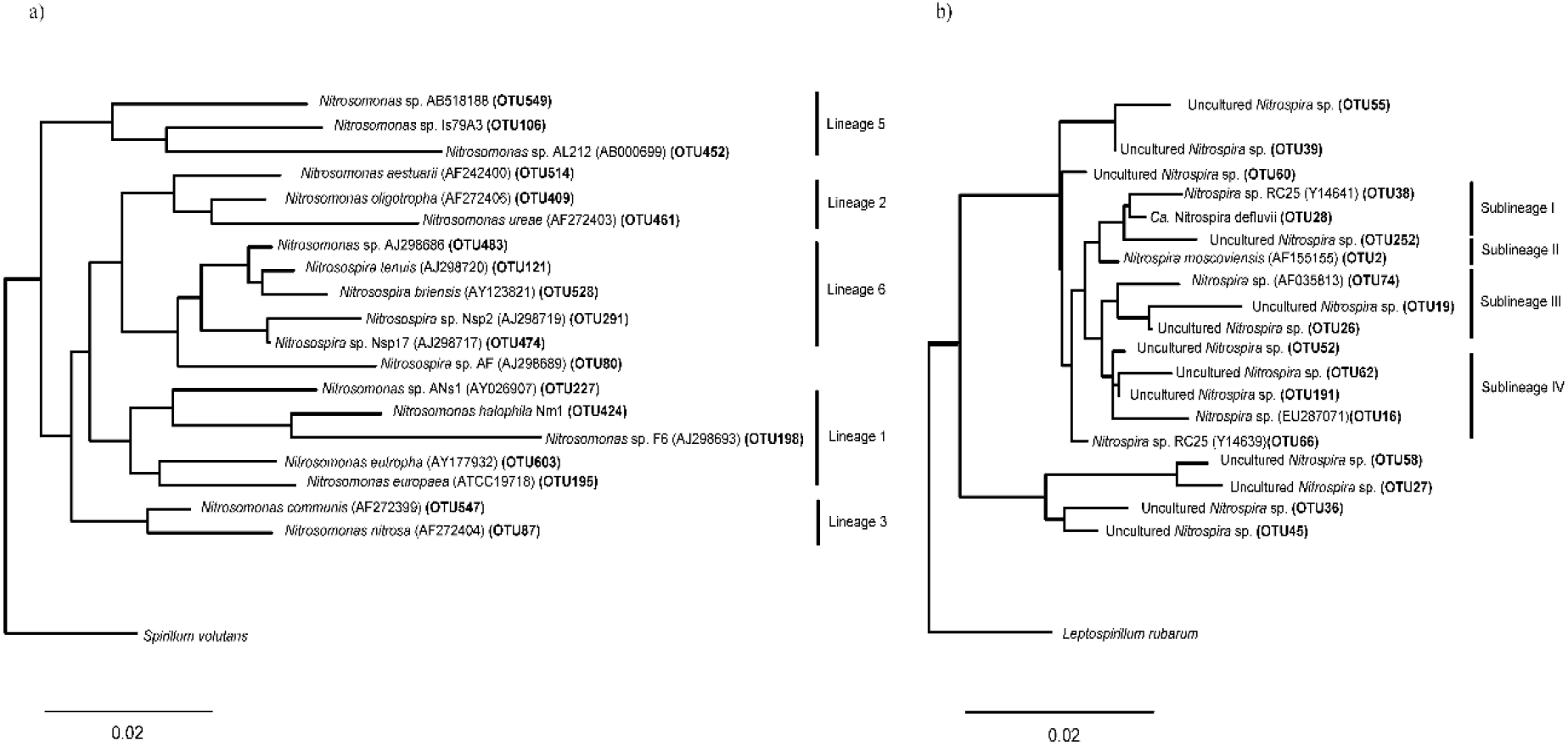
Dendograms showing phylogenetic positions of most dominant nitrifiers at species level based on sequences obtained after amplification of *amoA* and *Nitrospira*-related *nxrB* genes in activated sludge of Test Reactor and influent seed. Phylogenetic trees for most dominant AOB (a) and NOB (b) populations, identified based on computed Simpson diversity numbers representing number of dominant species in communities. Bracketed OTU numbers highlighted in bold belong to the most dominant AOB and *Nitrospira*-related NOB species detected in the samples. Entries without species designations are reported by their GenBank accession numbers. The scale bars represent 0.02 substitution per nucleotide position for the AOB and NOB phylogenetic trees.

**Fig. S5.**
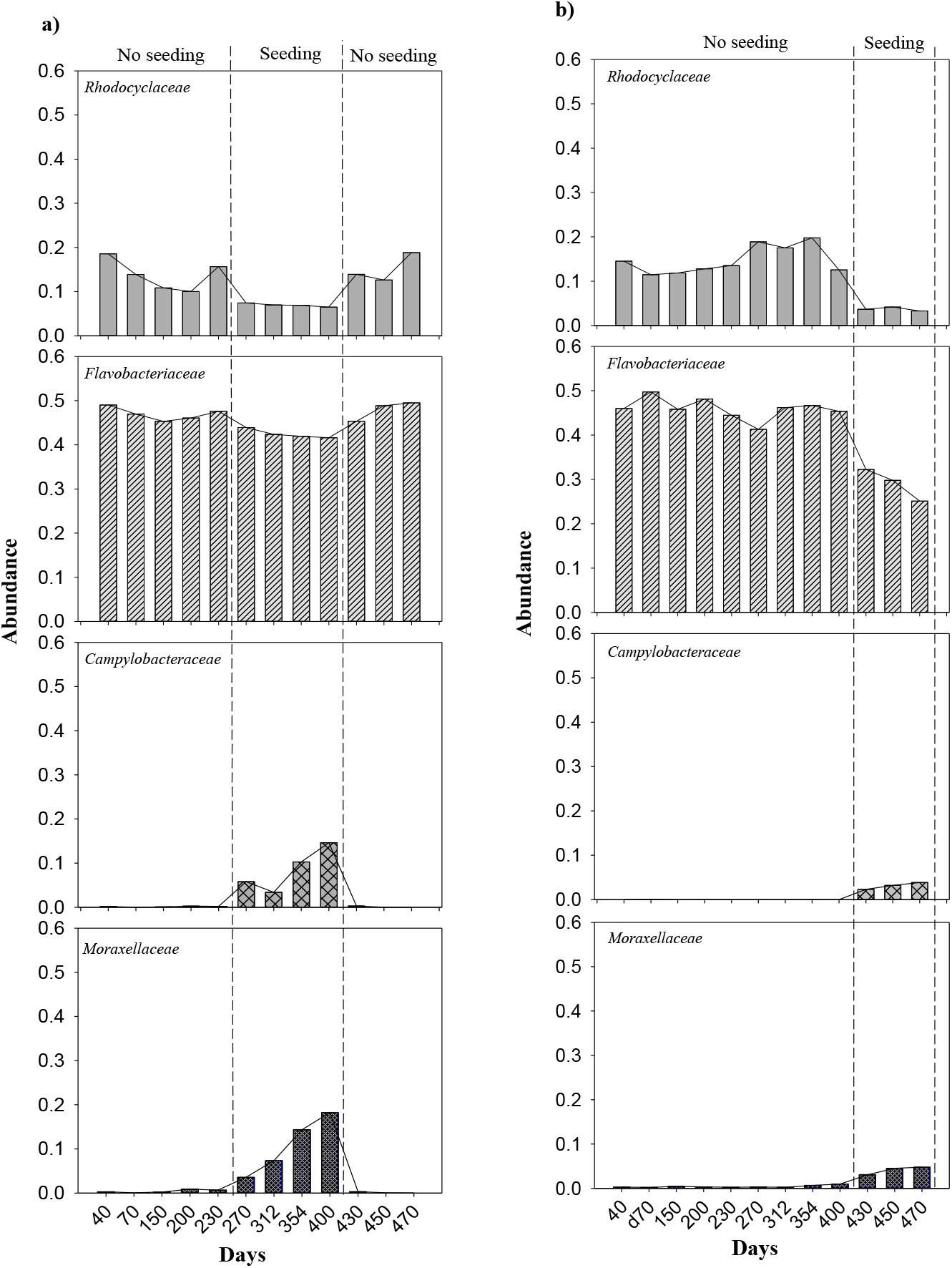
Abundance charts for the family *Rhodocyclaceae, Flavobacteriaceae, Campylobacteraceae* and *Moraxellaceae* for the Test Reactor (a) and Negative Control reactor (b). The seeding/no seeding periods are demarcated by the dashed lines.

**Fig. S6.**
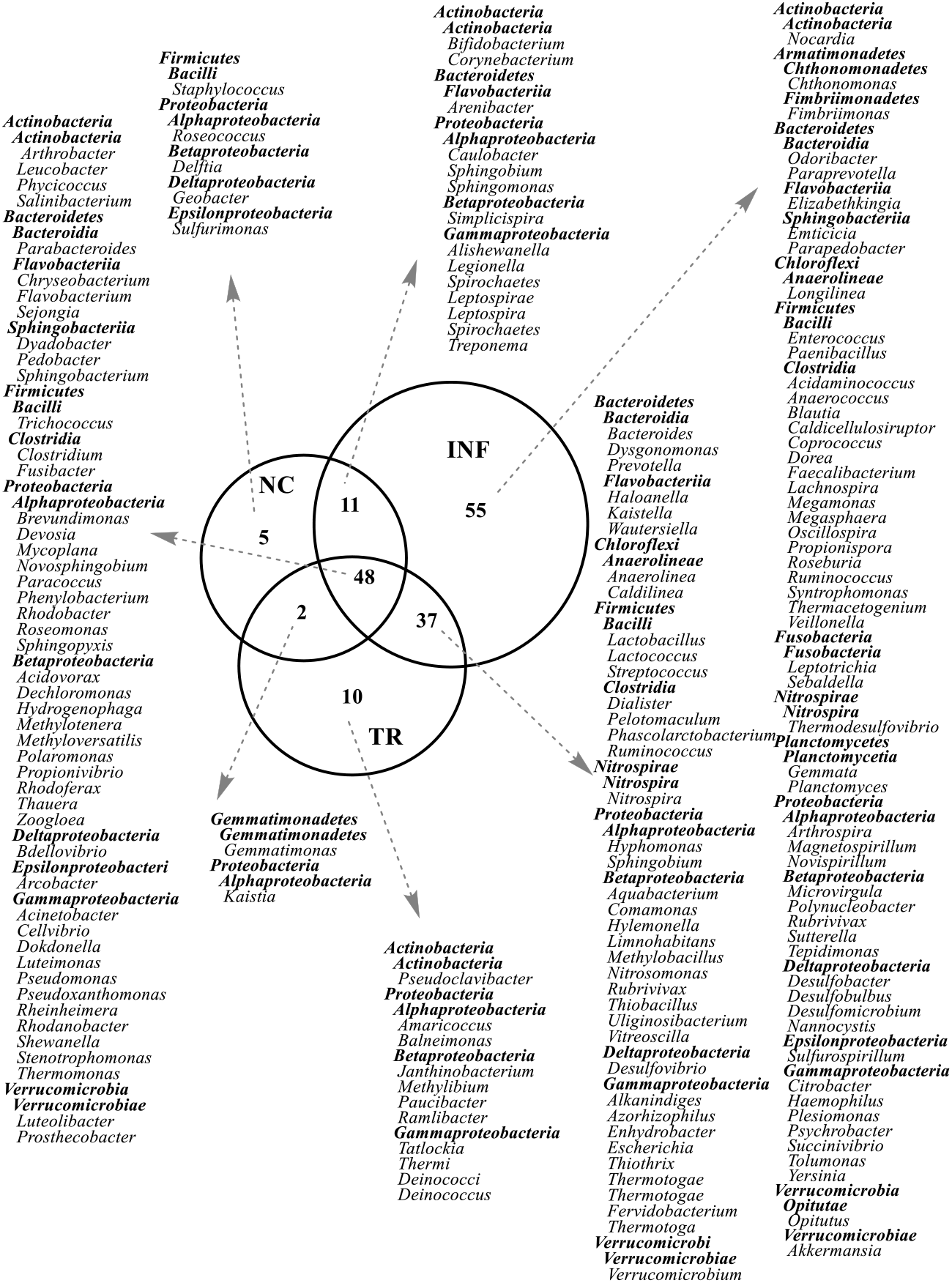
Venn diagram showing shared genera (indented with light font) between influent (INF), Test Reactor (TR) and Negative Control (NC) during period Day 270-400. Respective *phyla* (bold and not indented) and classes (bold and indented) to which the identified genera belong are also indicated.

